# Correlating physicochemical and biological properties to define critical quality attributes of a recombinant AAV vaccine candidate

**DOI:** 10.1101/2023.03.10.532114

**Authors:** Prashant Kumar, Michael Wang, Ozan S. Kumru, John Hickey, Julio Sanmiguel, Nerea Zabaleta, Luk H. Vandenberghe, Sangeeta B. Joshi, David B. Volkin

## Abstract

Recombinant adeno-associated viruses (rAAVs) are a preferred vector system in clinical gene transfer. A fundamental challenge to formulate and deliver rAAVs as stable and efficacious vaccines is to elucidate interrelationships between the vector’s physicochemical properties and biological potency. To this end, we evaluated an rAAV-based COVID-19 vaccine candidate which encodes the Spike antigen (AC3) and is produced by an industrially-compatible process. First, state-of-the-art analytical techniques were employed to determine key structural attributes of AC3 including primary and higher-order structures, particle size, empty/full capsid ratios, aggregates and multi-step thermal degradation pathway analysis. Next, several quantitative potency measures for AC3 were implemented and data were correlated with the physicochemical analyses on thermal-stressed and control samples. Results demonstrate links between decreasing AC3 physical stability profiles, *in vitro* transduction efficiency in a cell-based assay, and importantly, *in vivo* immunogenicity in a mouse model. These findings are discussed in the general context of future development of rAAV-based vaccines candidates as well as specifically for the rAAV vaccine application under study.

## Introduction

COVID-19, the disease caused by SARS-CoV-2, was declared a global pandemic by the World Health Organization (WHO) in March 2020 ^1,2^. Today, the disease accounts for over 500 million infections, 6 million deaths, and ongoing wide-scale damage to the global economy ^1,3^. To address the urgent need to save lives and mitigate these devastating effects, first-generation COVID-19 vaccines were developed and implemented at unprecedented speed ^3,4^ employing new vaccine platforms including adenovirus-based (AdV) vectors (e.g., Johnson & Johnson, University of Oxford-AstraZeneca) and mRNA-lipid nanoparticles (e.g., Pfizer-BioNTech, Moderna). First-generation COVID-19 vaccines are highly efficacious in protecting against severe disease and hospitalization caused by SARS-CoV-2 infections and have maintained efficacy against all variants identified to date ^5,6^. Recent mathematical modelling by Watson *et al.* estimated COVID-19 vaccines prevented ∼19.8 million deaths globally (63% global reduction in the deaths) between Dec 2020 to Dec 2021 ^7^. An additional 45-111% of deaths in low- and middle-income countries (LMICs) could have been averted if the global vaccination coverage target of 20 and 40%, as set by COVAX and WHO, respectively, had been met by each country by 2021 ^7^.

Although highly effective in protecting against severe illness, hospitalization and death ^8,9^, the new COVID-19 vaccines have not been adopted globally, largely due to high production costs, reliance on the cold-chain, economic inequity, and hesitancy ^10–15^. First-generation vaccines also display waning immunity and require multiple boosters to retain protective responses ^8,13,16,17^. One of the major cost drivers for COVID-19 vaccines is the requirement for cold and ultra-cold chain for vaccine production and distribution ^18^. For example, mRNA-based vaccines must be stored frozen at temperatures between -15 to -80°C to maintain potency^18^; together with a requirement for multiple doses, this significantly increases the overall cost ^18,19^. Viral vector platforms consisting of non-enveloped viruses such as AdV and rAAV based-vaccines, are relatively more thermostable if properly formulated ^20,21^ (e.g., AdV-based COVID-19 vaccine can be stored at 2-8°C)^22^. Another limitation of currently available COVID-19 vaccines is their administration by injection, which must be performed by a trained medical professional and is further complicated by the need for multiple doses for full immunization including booster doses ^19,23,24^. The WHO has outlined a target product profile for COVID-19 vaccines that would greatly enhance global access, especially in low- and middle income countries (LMICs), including a thermostable vaccine administered via a single primer dose followed by infrequent boosters ^25^.

In the past decade, recombinant adeno-associated virus (AAV) vectors have been proven safe and effective agents for gene therapy with three approved products and many ongoing clinical trials (see Discussion). Their utility as vaccine carriers has been studied less; prior to COVID-19, several studies in mice, ferrets, and monkeys indicate their ability to induce potent cellular and humoral immunogenicity to various vaccine targets in infectious disease and cancer ^26–33^. In humans, several studies in HIV showed overall safety yet variable immunogenicity ^34–36^. Clinically, rAAV vectors offer many potential advantages as a vaccine platform, both to protect against a wide variety of infectious diseases, as well as to specifically address the limitations of first-generation COVID-19 vaccines ^35,37–39^. Recently, a rAAV-based COVID-19 vaccine candidate AAVCOVID19-3 (referred to herein as AC3) has shown great promise in pre-clinical animal studies ^38,40^. AC3 is composed of the AAVrh32.22 capsid, which has been shown to have a unique pro-inflammatory activity among AAVs ^41–43^, and a recombinant genome containing inverted terminal repeats from AAV2 and genetic sequences encoding for the S1 subunit of the Wuhan S protein ^38^. AAVrh32.22 is derived from a Rhesus AAV isolates and is phylogenetically different from naturally circulating AAV capsid serotypes in humans, as well as others in use or under development for human gene therapy ^31,38,44^. A recent study by Zabaleta et al, demonstrated a single-injection of a thermally stable AC3 provided durable neutralizing immune responses and protection against SARS-CoV-2 infection in mouse and non-human primates ^38^.

As commercial products and clinical candidates in gene therapy, rAAV vectors are commonly formulated as frozen liquids stored at -80°C for administration by injection into site-specific targets. For use as vaccines to improve global access (see Discussion), rAAV vectors will need to be stabilized as low-cost, liquid formulations for storage in the refrigerator (or potentially at room temperature) and administered by routine IM injection (or potentially orally administered). Preliminary studies indicate AC3 to retain immunogenicity after storage at room temperature for 1 month ^38^. In this work, we performed a series of analytical characterization, accelerated stability and mouse immunogenicity studies with AC3 as a case study to enable future vaccine development of rAAV vectors. By determining the inter-relationships between physicochemical properties, *in vitro* transduction efficiency and *in vivo* performance in mice, these findings identify critical quality attributes (CQAs) of this AC3 vector as a vaccine candidate and provide a case study to further establish the well-characterized nature of rAAV vectors for use as a potential low-cost, stable next-generation vaccine platform.

## Results

As outlined in detail below, we first employed a suite of analytical characterization tools to quantify and comprehensively characterize the physicochemical properties of AC3 vector. Second, we elucidated the thermal inactivation mechanism of this rAAV vector using a subset of these methods. We then developed a cell-based *in vitro* transduction efficiency assay, analyzed thermally-stressed AC3 samples, and correlated *in vitro* potency results with stability-indicating physical and genome copy assays. Finally, we evaluated AC3 samples with varying alterations in their physicochemical and transduction efficiency properties for effects on *in vivo* performance as a vaccine candidate in a mouse immunogenicity assay.

### Physicochemical characterization of AC3 vector

The structural integrity of the AC3 capsid proteins (viral proteins VP1-3, see **Figure 1**) were determined using a combination of sodium dodecyl-sulfate polyacrylamide gel electrophoresis (SDS-PAGE), capillary electrophoresis sodium dodecyl sulfate (CE-SDS) and liquid chromatography-mass spectrometry (LC-MS) intact and peptide mapping analyses. First, SDS-PAGE (**Figure 1A**) showed distinct bands corresponding to the VP1, VP2 and VP3 proteins migrating at the expected ∼80, ∼65 and ∼60 kDa, respectively. The relative abundance ranges of the VP1:VP2:VP3 proteins were (7-10%):(16-21%):(72-74%) leading to an approximate relative ratio of 1:2:9. Similar results were observed under reducing conditions indicating no disulfide linkages in these proteins. CE-SDS analysis (**Figure 1B**) provided essentially equivalent results under both non-reduced and reduced conditions, but additionally allowed for quantification of VP3 protein truncations present in a range of 11-13% and 16-18% under non-reducing and reducing conditions, respectively.

**Figure 1:**
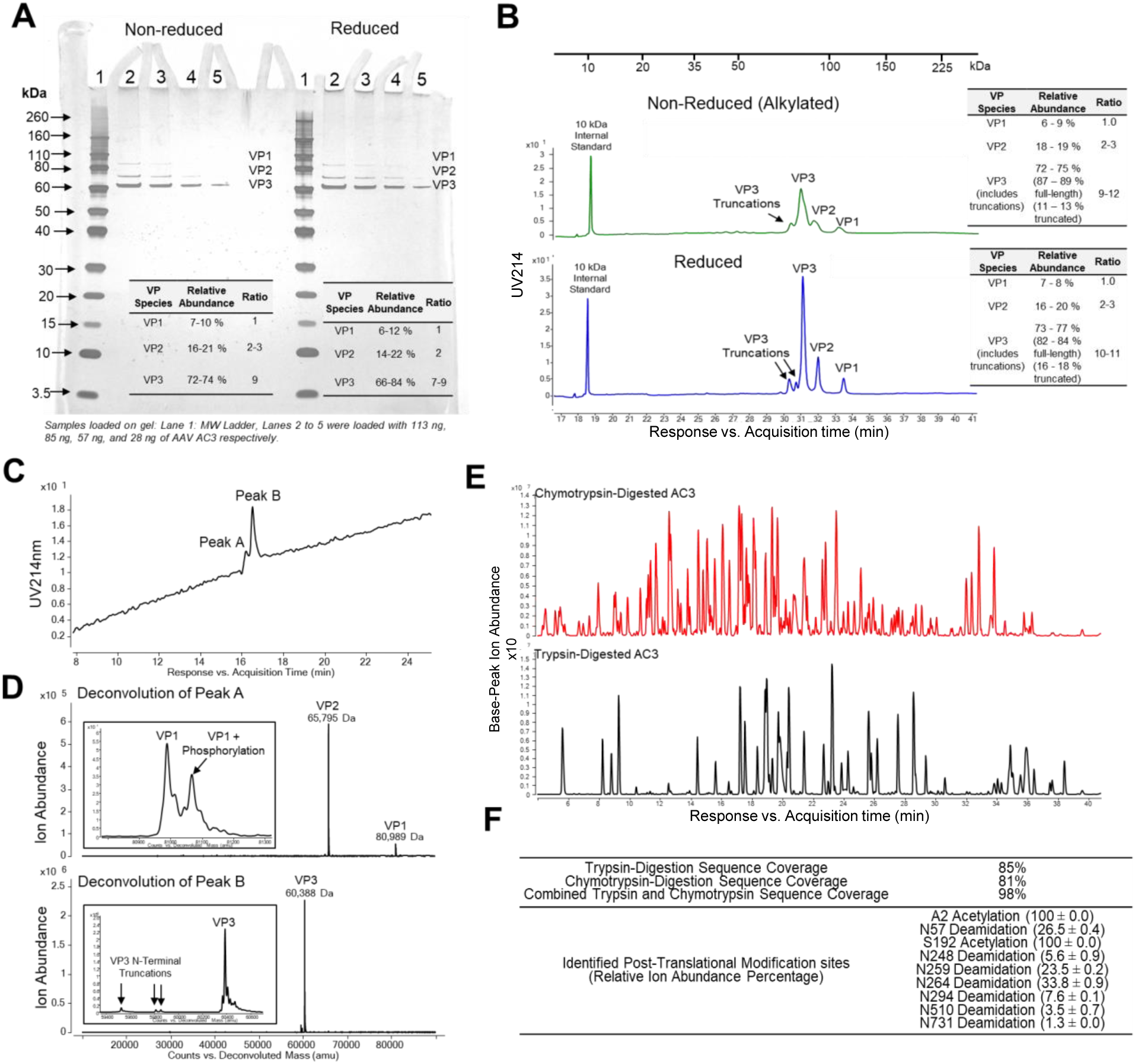
Primary structure and post-translational modification analysis of AC3 capsid proteins VP1, VP2 and VP3. Molecular weight and relative abundance of VP1, VP2 and VP3 as measured by (A) non-reducing and reducing SDS-PAGE, or (B) non-reducing and reducing CE-SDS. Intact LC-MS analysis of AC3 capsid proteins including (C) RP-UPLC of the non-reduced AC3 construct, and (D) Intact Mass analysis of the major VP species comprising the RP-UHPLC Peaks A and B. Minor species near the VP1 or VP3 proteins were enlarged for easier visualization. LC-MS peptide mapping analysis of AC3 vector including (E) representative base-peak ion chromatograms of chymotrypsin- (red trace) or trypsin-digested (black trace), and (F) trypsin and chymotrypsin peptide maps coverage and identified post-transitional modifications.

Additional post-translational modifications (PTMs) were identified using two different LC-MS techniques. During chromatographic separation on a reversed-phase column, two peaks were observed with VP1 and VP2 proteins co-eluting (Peak A) followed by the VP3 protein (Peak B) (**Figure 1C**). Subsequent intact mass spectrometry analysis of Peak A showed VP2 as a single species (65,795 ± 1 Da) while VP1 was two species with PTMs including an N-terminal acetylated species with (81,067 ± 2 Da) or without a single phosphorylation (80,988 ± 1 Da) (**Figure 1D**). LC-MS peptide mapping provided site-specific information about the PTMs within a combined sequence coverage (both trypsin, chymotrypsin digestions) of 98% (**Figures 1E and Supplemental S2**). For the VP1 protein, PTMs included the absence of M1 and an N-terminal acetylation at A2, while multiple VP3 species were observed. The mass of the major VP3 species (60,388 ± 0 Da) was consistent with the expected mass (residues 192-733) containing an N-terminal acetylation. Two low abundant VP3 variants were also observed that corresponded to N-terminal truncations (residues 197-733 and 199-733). In summary, a series of PTMs were identified including truncations of VP3, phosphorylation of VP1, two N-terminal acetylation sites at specific Ala and Ser residues in VP1 and VP3, and finally multiple Asn deamidation sites across the entire sequence (**Figure 1F**).

Characterization of the assembled AC3 capsids (including particle size, morphology, ratio of empty-full capsids and number of genome copies) was performed using a combination of dynamic light scattering (DLS), transmission electron microscopy (TEM), sedimentation velocity analytical ultracentrifugation (SV-AUC) and a genome titration assay, respectively. The hydrodynamic diameter of AC3 capsid was 31-33 nm (± 2 nm), depending on the calculation method used for DLS analysis (**Figure 2A**). The AC3 particles displayed a spherical morphology with a diameter of ∼26 ± 2 nm by TEM analysis (**Figure 2B**). These small differences in size range are attributed to differences in analytical principles including analysis of dried (TEM) vs solution (DLS) AC3 samples ^46,47^. TEM using negative staining with uranyl acetate showed 36 ± 3% full capsids were present in the AC3 sample (**Figure 2B**) but was unable to distinguish empty vs. partially full capsids. SV-AUC analysis, however, better separated multiple AC3 species of varying sedimentation coefficients which were classified as full capsids (90-110 S), partially-filled capsids (60-90 S), and empty capsids (50-60 S) (**Figure 2C**). A few large agglomerates (> 110 S) and other product or process related impurities (< 50S) were also present. The AC3 preparation comprised of 37 ± 9% full, 39 ± 3% partial, and 14 ± 6% empty capsids with low amounts of agglomerates and impurity/fragments (1 ± 1% and 8 ± 4%, respectively). Finally, the total genome copy numbers in the AC3 preparation were determined (∼2E+12 gc/mL) and were shown to be independent of DNase-I treatment (**Figure 2D**), a result indicating high-purity with absence of DNA outside AAV capsid.

**Figure 2:**
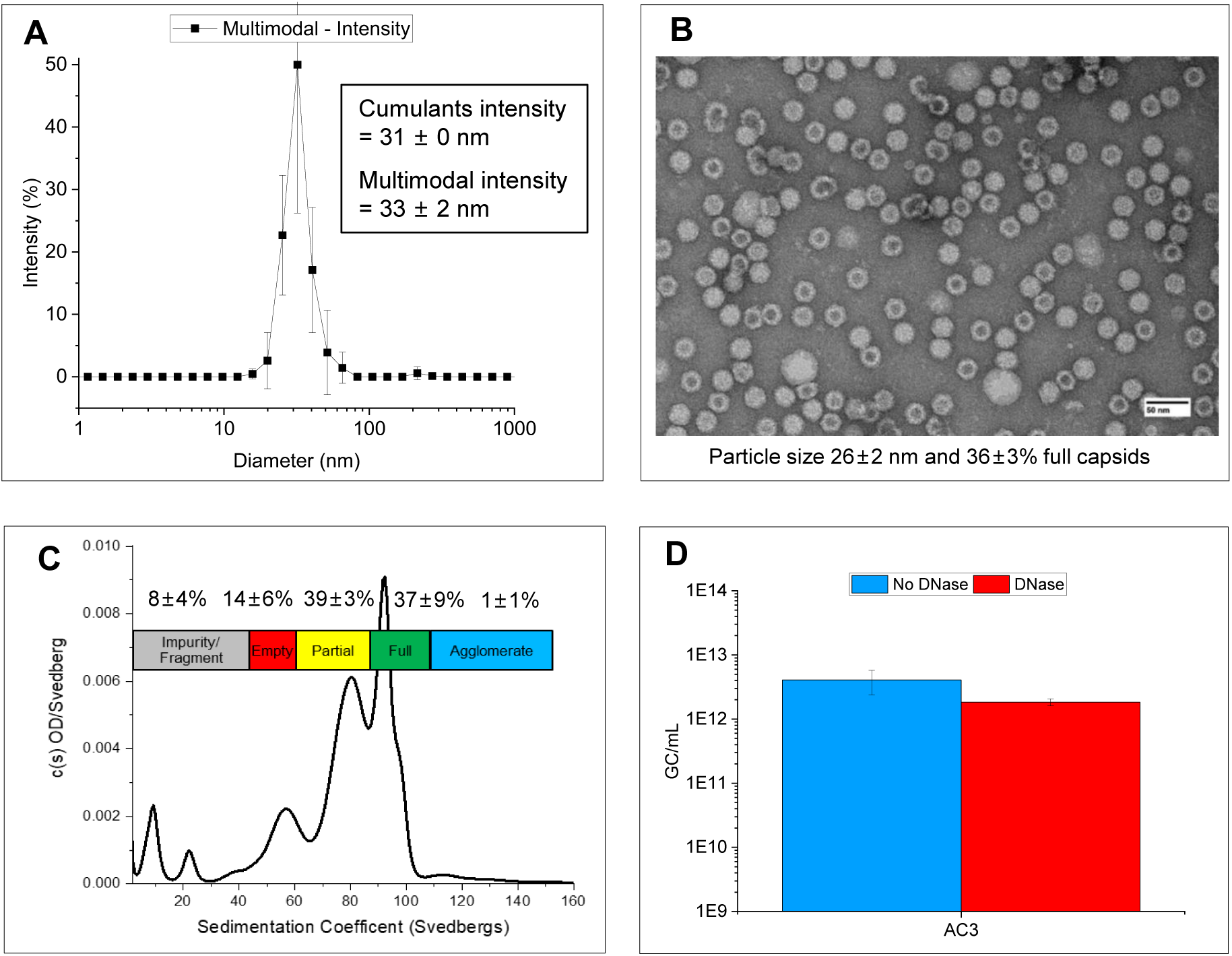
Characterization of the size, morphology, empty-full ratios and genome copies of AC3 vector. (A) Hydrodynamic diameter measured by DLS and fit using cumulants intensity and multimodal intensity analysis, (B) representative TEM image showing size, morphology and percent full capsid species, (C) SV-AUC profile displaying AC3 various species based on sedimentation coefficient ranges, (D) Total genome copy numbers of non-DNase-treated and DNase-treated AC3 measured using genome titration assay. All samples were analyzed at n ≥ 3 and error bars represent 1 SD.

### Mechanism of Thermal Degradation of AC3 vector

As an initial evaluation of the physical stability of the AC3 vector, both unstressed and heat-stressed (60°C 10 min) samples were analyzed by SV-AUC and TEM. SV-AUC sedimentation coefficient distributions for unstressed and thermal-stressed AC3 (**Figure 3A**) indicated a large loss of full and partially-full capsids and a relative increase in empty capsids and fragments in the thermal-stressed AC3 as determined by total peak areas (**Figure 3B**). TEM analysis of the same unstressed and thermal-stressed AC3 samples (**Figure 3C-D**) showed a large reduction in the proportion of full AC3 capsids (36 ± 3% vs 13 ± 1%, respectively). Interestingly, no significant increase in particle size of AC3 was noted in unstressed (26 ± 2 nm) and thermal-stressed (28 ± 1 nm) by TEM, however, a trend of more agglomerates of individual AC3 particles present in the thermally stressed sample was visually observed (**Figure 3C-D**). The agglomerates observed by TEM were likely not observed by SV-AUC due to their immediate sedimentation during analysis. In summary, heat-treatment of AC3 led to large losses in full and partially-full capsids with a concomitant increase in empty capsids, fragments and agglomerates.

**Figure 3:**
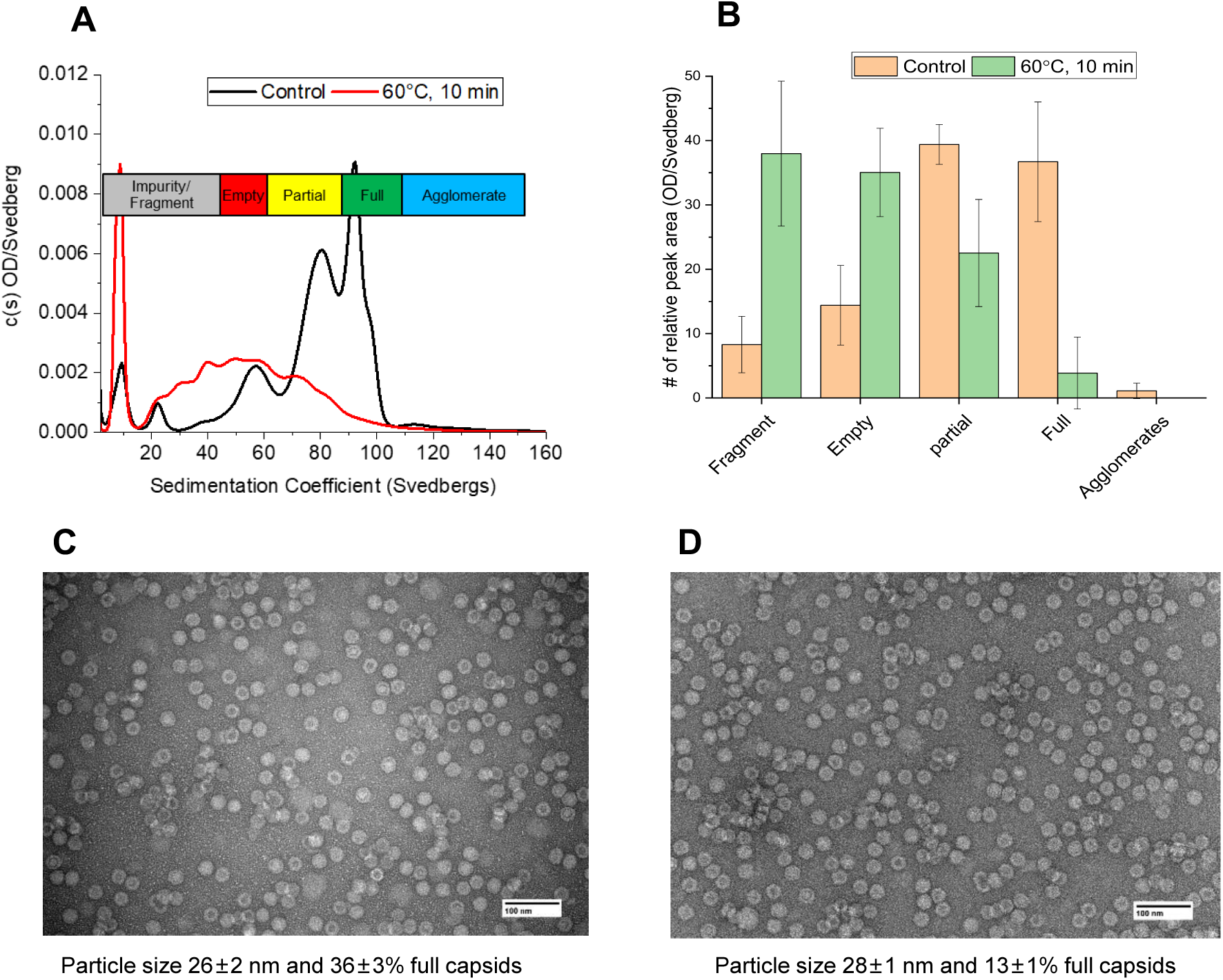
Effect of thermal stress treatment of AC3 capsid on vector particle size, morphology and empty-full capsid ratio. Stressed samples were incubated for 10 min at 60°C and then compared to unstressed samples by (A) SV-AUC analysis showing sedimentation coefficient distributions, and (B) SV-AUC total peak areas of each species from triplicate analyses (n = 3 ± 1 SD). (C, D) TEM analysis showing representative unstressed (C) or thermal-stressed (D) samples. A total of 250 particles per AC3 sample were used to calculate the particle size and relative abundance of full capsids.

To better understand the mechanistic aspects of this observed thermal degradation of AC3 vector, we employed temperature ramp studies coupled to various fluorescence spectroscopy detection methods (both intrinsic tryptophan and extrinsic dyes) ^48,49^, as well as light scattering methods (both dynamic and static) to evaluate structural alterations to the AC3 capsid (**Figure 4A-E**). Three major structural transitions were observed as a function of increasing temperature: a “small” transition range (∼35° to 60°C), a “medium” transition range (∼55 to 65°C) and a “major” transition range (∼64 to 70°C) (**Figure 4A-E**). Cartoon representations of three these structural transitions for each of AC3 species (i.e., full, partially-filled, and empty capsids) are displayed (**Figure 4F**). The “small” transition (∼35 to 60°C) displays minor structural changes in the VP1-3 proteins of the AC3 capsid as observed by increasing signals in the intrinsic (Trp) and extrinsic (SYPRO orange dye) fluorescence measurements. The increase in maximum peak position for intrinsic Trp fluorescence demonstrates the average Trp residue is exposed to a more hydrophilic (aqueous) environment, while the increase signal of SYPRO orange dye shows exposure to a more hydrophobic milieu (binds to protein). The SYBR Gold nucleic acid dye and light scattering methods show no increase in signal, a result indicating the dye could not access the nucleic acid inside the AC3 capsid, and no capsid agglomeration, respectively.

**Figure 4.**
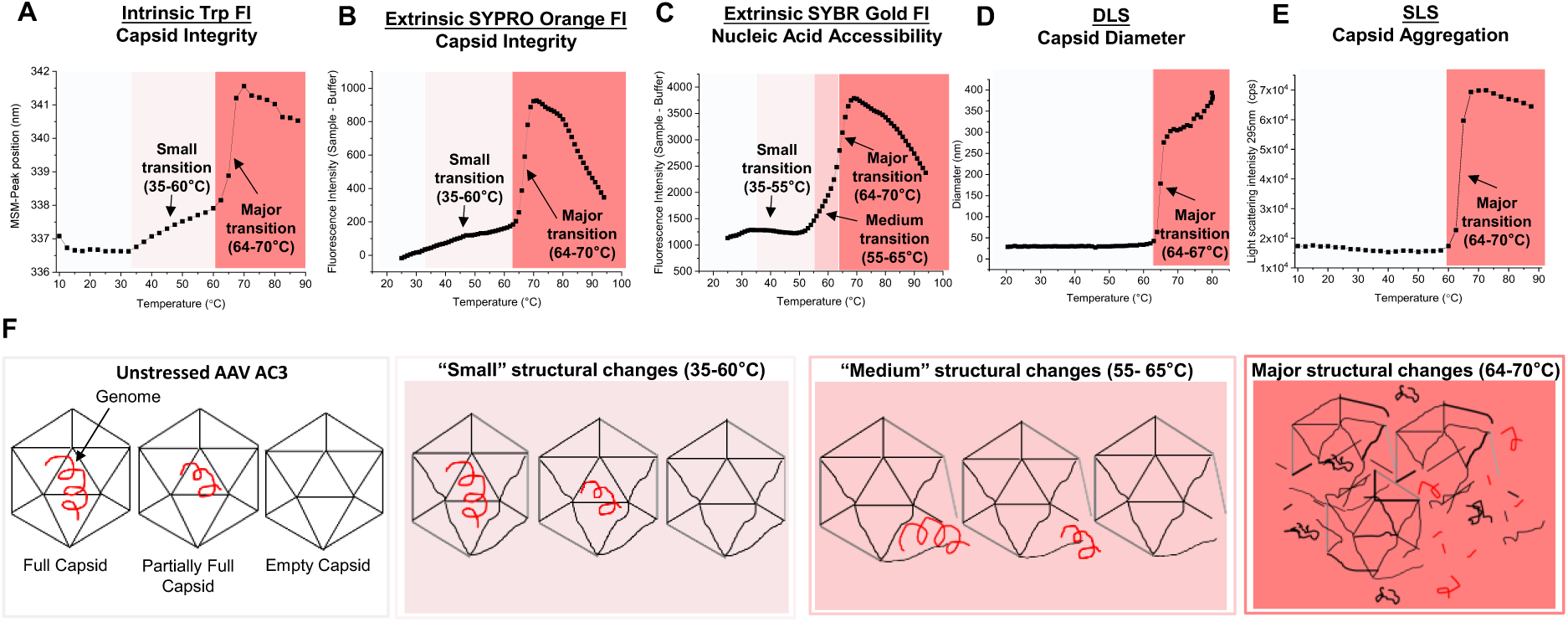
Mechanistic studies on thermal degradation of AC3 capsid as measured by a combination of fluorescence spectroscopy and light scattering studies. As the temperature of the AC3 containing solution was ramped up to 90°C, vector degradation was monitored by a combination of (A) intrinsic fluorescence, (B, C) extrinsic fluorescence in the presence of SYPRO orange and SYBR gold dyes, (D) dynamic light scattering, and (E) static light scattering. Readouts from the analytical methods are grouped by temperature ranges showing “small” (35-60°C), “medium“ (55-65°C) and “major” (64-70°C) structural transitions of AC3 vector. (F) Cartoon representation of different AC3 species in the purified sample (full, partially-full, empty capsids) and associated structural alteration events in the temperature ranges for the “small”, “medium” and “major” transitions.

More dramatic structural alterations in the AC3 capsid are observed as the temperature increases further. In the “medium” transition temperature range (∼55 to 65°C), not only do the intrinsic (Trp) and extrinsic (SYPRO orange dye) fluorescence measurements continue to increase, but a concomitant increase in SYBR Gold fluorescence is seen initiating at ∼55°C. This result indicates moderate structural alterations in the AC3 capsid resulting in release of nucleic acid. For the “major” transition temperature range (64 to 70°C), the AC3 capsid undergoes much more substantial degradation including protein denaturation (i.e., exposure of interior, hydrophobic regions of the viral proteins indicated by intrinsic and extrinsic fluorescence), nucleic acid accessibility (SYBR Gold), and capsid aggregation (DLS and static light scattering (SLS)). With this mechanistic understanding of the temperature induced physical degradation of AC3 vector, we then sought to determine how such structural alterations can affect the biological activity of AC3, both in terms of gene transduction as well as a vaccine immunogen, as described below.

### Cell-based transduction efficiency assay and evaluation of AC3 vector before and after thermal stress

Genome copy number of rAAV is routinely used for dosing of rAAV but it fails to fully capture the biological activity of the AAV preparations. Thus, animal models are typically used to test potency of rAAV preparations, however, animal studies are expensive, low-throughput and time consuming and are thus impractical for process and formulation development work. We therefore developed a high-throughput real-time quantitative polymerase chain reaction (RT-qPCR) based gene transduction assay for probing relative gene expression of the AC3 vector. The principle of transduction efficiency method (i.e., *in vitro* potency) is based on detection and amplification of mature mRNA expressed by the vector using a specific TaqMan™ probe in a two-step RT-qPCR assay. The probe is specific for mature transgene mRNA (after splicing) and does not recognize the precursor transgene mRNA (prior to splicing) or recombinant vector DNA (**Figure 5A**). The presence of spliced mRNA product was validated by Phusion PCR and sequencing, and validation of assay specificity was performed using RT-qPCR in the presence or absence of reverse transcriptase and was found suitable for analysis of AC3 expression (**Supplemental Figure S1**). A range of estimated MOI for AC3 was evaluated and the linear range of the assay was found to be 2-4.5 log estimated MOI with an R-squared value of 0.99 (**Figure 5B**).

**Figure 5:**
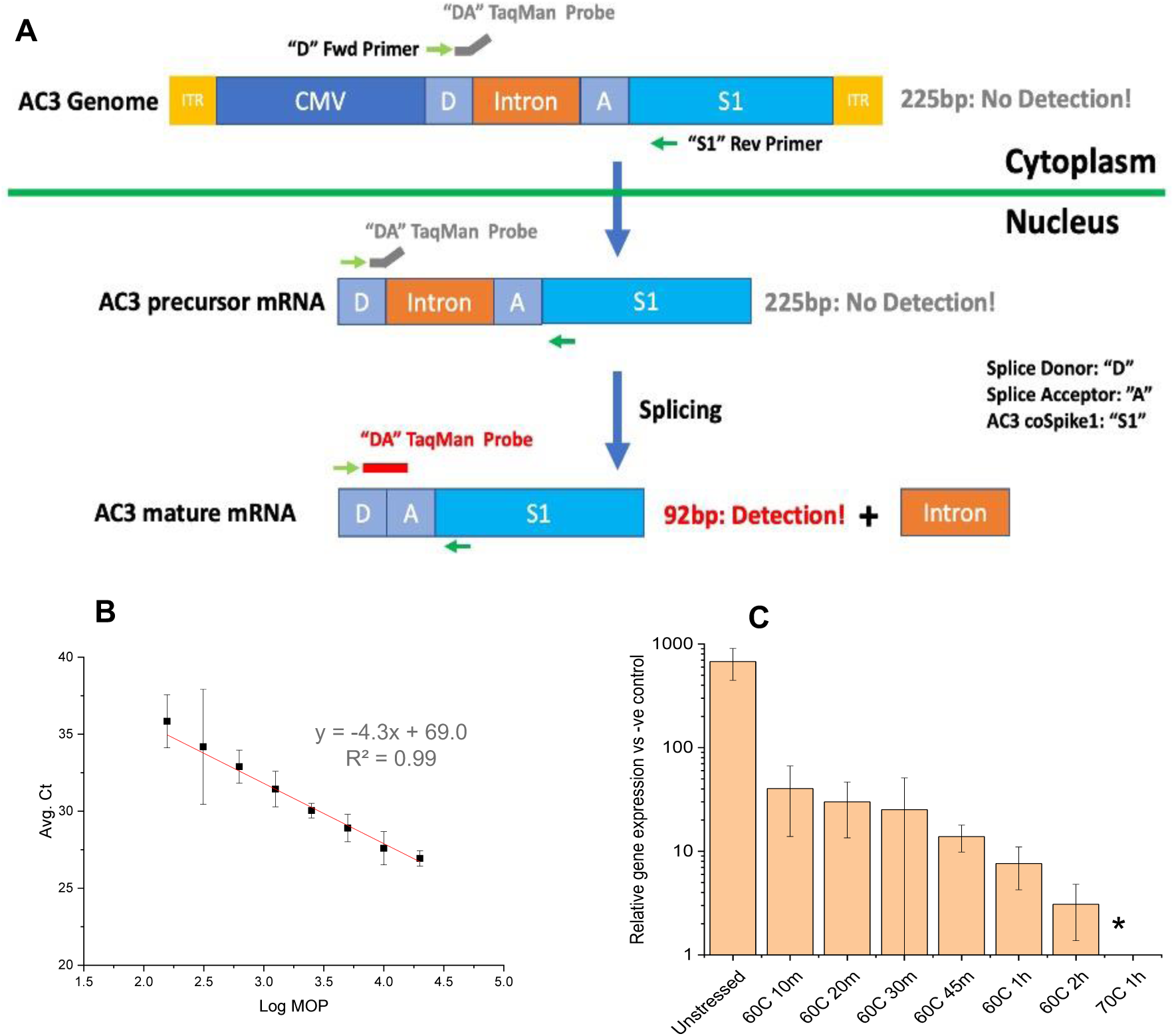
*In vitro* transduction efficiency assay design and infectious titer results with AC3 samples before and after thermal stress. (A) Assay design and (B) linear range of the assay (Ct vs Log MOP). (C) The relative gene expression of AC3 vector before (unstressed) or after heat treatment (10-120 min at 60°C or 60 min at 70°C). All AC3 samples were analyzed in quadruplicate and error bars represent 1 SD. An asterisk (*) denotes the relative gene expression was below the limit of quantification. Ct-threshold cycle, MOP-multiplicity of particles, D-splice donor, A-splice acceptor, S1- AC3 coSpike1.

To establish selectivity and stability-indication of the *in vitro* potency assay of AC3, samples were incubated under two different conditions (i.e., from 10-120 min at 60°C, or for 60 min at 70°C) and then analyzed by RT-qPCR. A progressive reduction in relative gene expression with respect to negative control (wildtype Adeno 5 only, no AC3) was observed as a function of increasing thermal stress (**Figure 5C**). The results indicated an initial *in vitro* potency loss of ∼1 log (after 10 min at 60°C) which remained relatively unchanged (up to 30 min at 60°C), followed by a relatively slower loss in *in vitro* potency (up to 120 min at 60°C).

These same unstressed and thermally-stressed AC3 samples were also analyzed using the physical methods described above, and the results obtained were then correlated using Pearson’s method (i.e., the Pearson’s correlation coefficient denoted by r lies between -1 and 1 and its absolute value in the range of 0.1–0.3 indicates small, 0.3–0.5 medium and 0.5–1.0 large correlations ^50^) with infectious titers measured using the *in vitro* transduction efficiency assay. For example, after application of thermal stress, the reduction in genome copy numbers (in presence of DNase-1) correlated strongly with the reduction in gene expression (**Figures 6A, D**).

**Figure 6:**
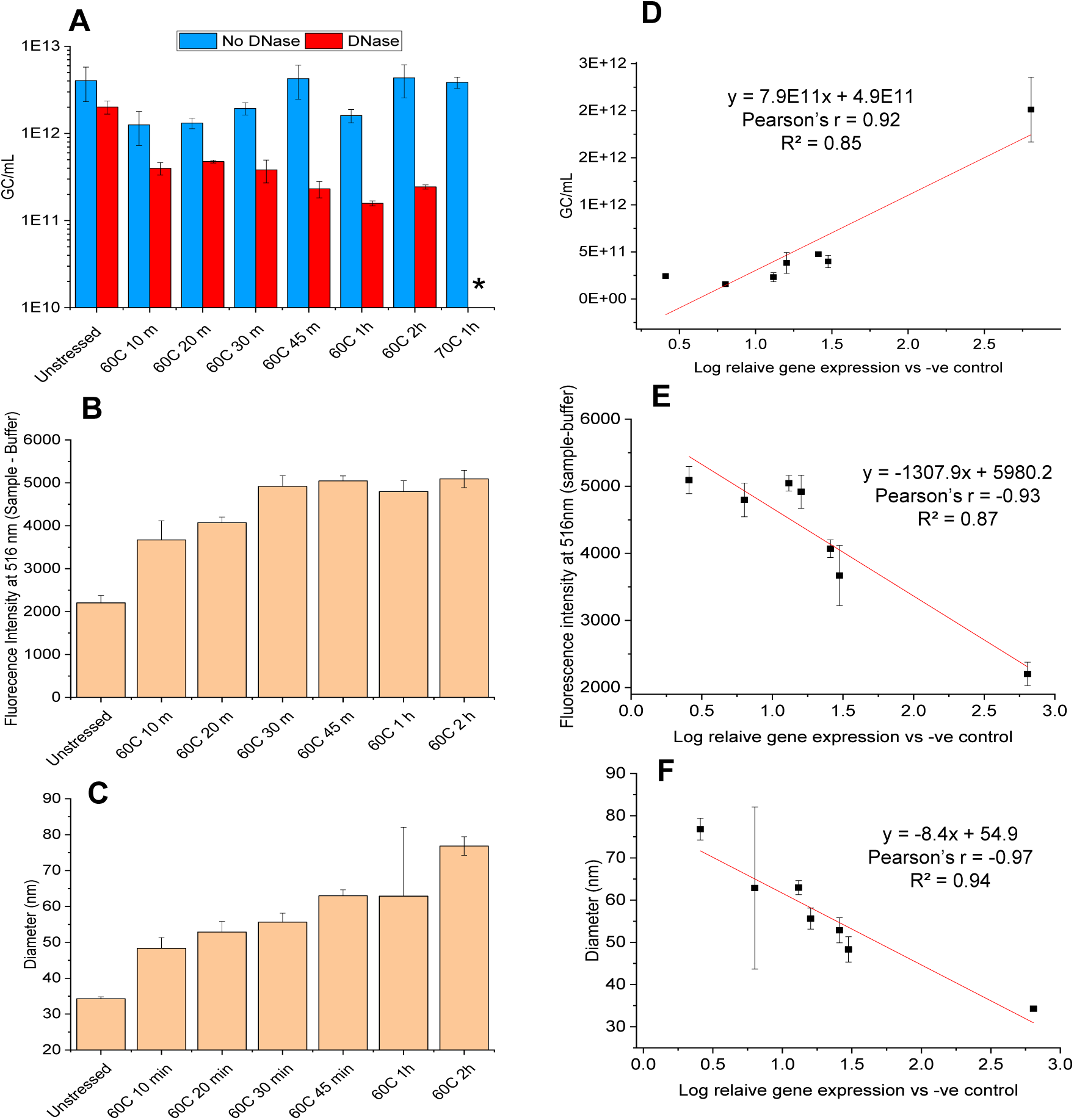
Analysis of unstressed and heat-stressed AC3 vector samples and correlation of results with infectious titers as measured by the *in vitro* transduction efficiency assay. (A) AC3 genome copy numbers as measured by genome titration assay, (B) Nucleic acid accessibility of AC3 evaluated by SYBR Gold fluorescence assay, and (C) mean particle diameter obtained by DLS multimodal intensity analysis. Correlation of results with measured *in vitro* transduction efficiency titers of same samples for (D) genome copy numbers, (E) SYBR Gold fluorescence intensity at 516nm, (F) mean particle diameter. Thermally stressed samples were incubated from 10-120 min at 60°C or 60 min at 70°C. All AC3 samples were analyzed at n ≥ 3 and error bars represent 1 SD. An asterisk (*) denotes the gc/mL number was below the limit of quantification.

When analyzed in absence of DNase-I, no change in total genome copy number (and thus no such correlation) was observed indicating only encapsidated genome was responsible for infectivity. Nucleic acid accessibility probed SYBR Gold fluorescence analysis showed two-fold increase signal in stressed samples after thermal incubation (120 min at 60°C), showing higher amounts of solvent accessible nucleic acid and a good correlation with loss in *in vitro* transduction efficiency (**Figure 6B, E**). Finally, the mean hydrodynamic diameter of AC3 samples probed using DLS displayed an increase in particle size from ∼33 nm (unstressed) to ∼80 nm (120 min at 60°C) and a good correlation with loss in *in vitro* transduction efficiency (**Figure 6C, F**). Interestingly, changes in the signals as a function of thermal stress for other physical methods did not correlate well (Pearson’s r 0.57 to 0.77) with the *in vitro* transduction efficiency assay (**Supplemental Figure S3**). For example, evaluations of various incubated AC3 samples by (1) conformational stability (i.e., thermal melting temperatures or Tm values), (2) the extent of SYPRO orange binding, or (3) the level of sub-visible particles did not correlate with loss of *in vitro* potency. Interestingly, although the total number of sub-visible particles (2-100 microns) of unstressed and stressed AC3 (10 min at 60°C) indicated no notable change after thermal stress, the proportions of larger sub-visible particles (40-100 µM) increased (**Supplemental Figure S4**). In summary, three physical methods correlated well (Pearson’s r > 0.85) between their signal changes and losses observed in the *in vitro* transduction efficiency assay (for AC3 samples after incubation for 10-120 min at 60°C or 60 min at 70°C). These included the genome copy, differential scanning fluorimetry (DSF), SYBR Gold (nucleic acid accessibility) and DLS (capsid aggregation) assays.

Finally, to better understand the interrelationships between structural changes in the AC3 capsid (during thermal stress) and *in vitro* transduction activity, we repeated the thermal ramping experiments (as described in Fig 4) performed for mechanistic studies, but in this case without the fluorescent dye. We removed samples at specific temperatures (i.e., 40°, 50°, 60°, 70°C) and then measured genome copy numbers (**Figure 7A**) and *in vitro* transduction (**Figure 7B**). For the unstressed AC3 sample (4°C), comparable numbers of encapsidated and total genomes were observed, and these values were normalized to 100 percent for comparisons to stressed samples. Similarly, the gene expression values for the unstressed sample were determined and normalized to 100 percent. During the initial stages of temperature ramping (to either 40° or 50°C), no changes in encapsulated genomes or gene expression levels were observed in the AC3 samples.

**Figure 7:**
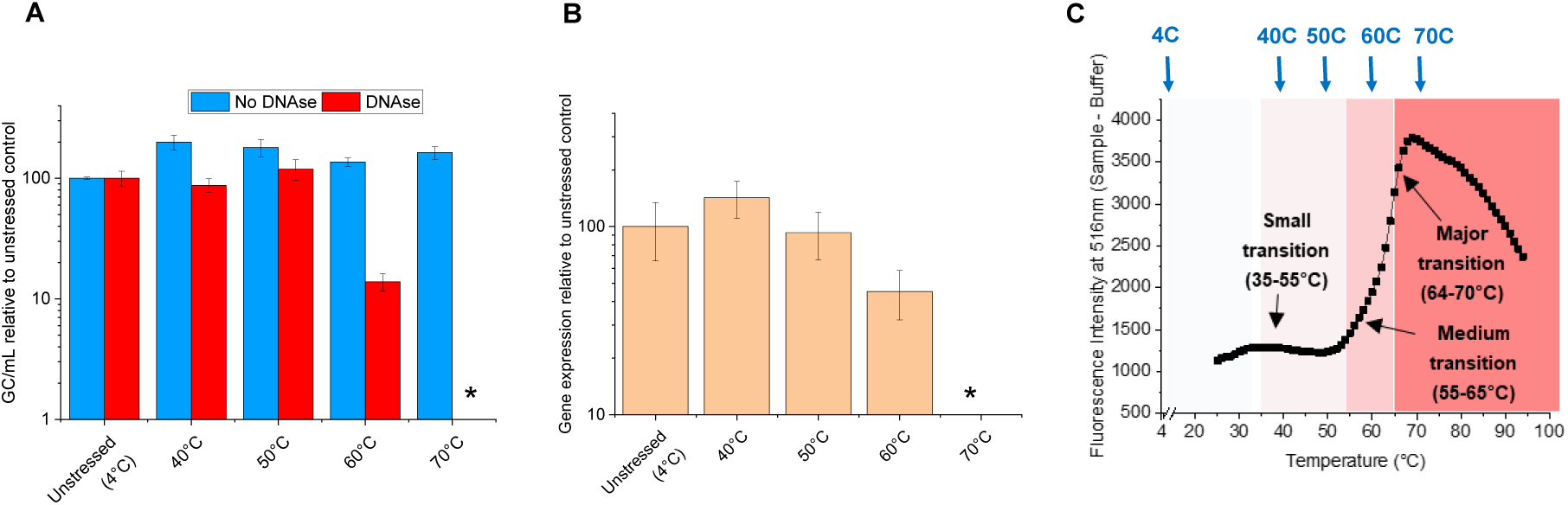
Relative genome copies and gene expression levels for unstressed vs. heat-stressed AC3 samples from temperature ramping studies. (A) relative genome copy numbers for heat-stressed AC3 samples (vs unstressed AC3 at 4°C) as measured by genome copy assay, (B) relative gene expression levels for heat-stressed AC3 samples (vs unstressed AC3 at 4°C) as measured by *in vitro* transduction efficiency assay. Samples were subjected to thermal ramping conditions identical to the SYBR Gold fluorescence assay results described in Figure 4 (reshown in here in panel C for ease of comparison), but with no dye added with samples taken when the temperature reached 40°, 50°, 60°, and 70°C. All samples were measured in quadruplicate and error bars represent 1 SD. An asterisk (*) denotes the gc/mL number was below the limit of quantification.

In contrast, for AC3 samples undergoing a temperature ramp to 60°C, notable partial losses in encapsidated genomes (∼80% loss) and gene expression levels (∼50-60% loss) were observed. When AC3 samples were ramped up to 70°C, a complete loss in both encapsidated genomes and gene expression was seen. Since we performed this thermal ramping experiment under the same conditions as the mechanistic studies (Figure 4), we could make additional direct comparisons (data reshown for SYBR Gold results in **Figure 7C** for ease of comparisons). During exposure to the “small” structural transition temperature range (∼35-55°C), nucleic acid remains inaccessible to SYBR Gold and there are no losses in encapsidated genomes and gene expression. During exposure to the “medium” structural transition temperature range (∼55-65°C), partial exposure of nucleic acid (SYBR Gold) correlated with partial loss of encapsidated genomes and gene expression. Finally, after exposure to the “major” structural transition temperature range (∼64-70°C), results demonstrate nucleic acid accessibility starts at ∼55°C, maximum losses of nucleic acid from the capsid corresponds to complete loss of encapsidated genomes and gene expression.

### Correlations between in vivo immunogenicity, in vitro transduction efficiency and physical stability profiles

We performed mouse immunogenicity studies with unstressed and stressed samples of AC3 characterized above by physicochemical and *in vitro* potency assays. AC3 samples were incubated (1-120 min at 60°C or 60 min at 70°C). The mouse immunogenicity study results were measured as geometric mean titers of antibody responses to the receptor-binding domain (RBD)^38^ in C57BL/6J mice 28 days after vaccination with 10E+11 gc of AC3. The same samples were also assayed for *in vitro* transduction efficiency. The total antibody response progressively decreased as a function of increasing thermal stress of the AC3 vector, and the trends agreed well with the *in vitro* transduction efficiency results (**Figures 8A**). A strong positive Pearson’s correlation (Pearson’s r = 0.99) was obtained for the measured potencies from both the assays i.e., the *in vivo* mouse immunogenicity and *in vitro* transduction efficiency assays (**Figure 8C**). Longitudinal antibody responses in mice treated with 10E+11 gc of AC3 before and after thermal stress were also measured (**Figure 8D**). The results indicate the highest geometric mean RBD IgG titers over time in the unstressed AC3 sample, and that decreased as a function of increasing thermal stress treatment of the AC3 vector at all measured time points. The RBD IgG titers were at their peak in the unstressed and all stressed AC3 samples at 21 days and remained unchanged up to the last measured time point of 56 days.

**Figure 8:**
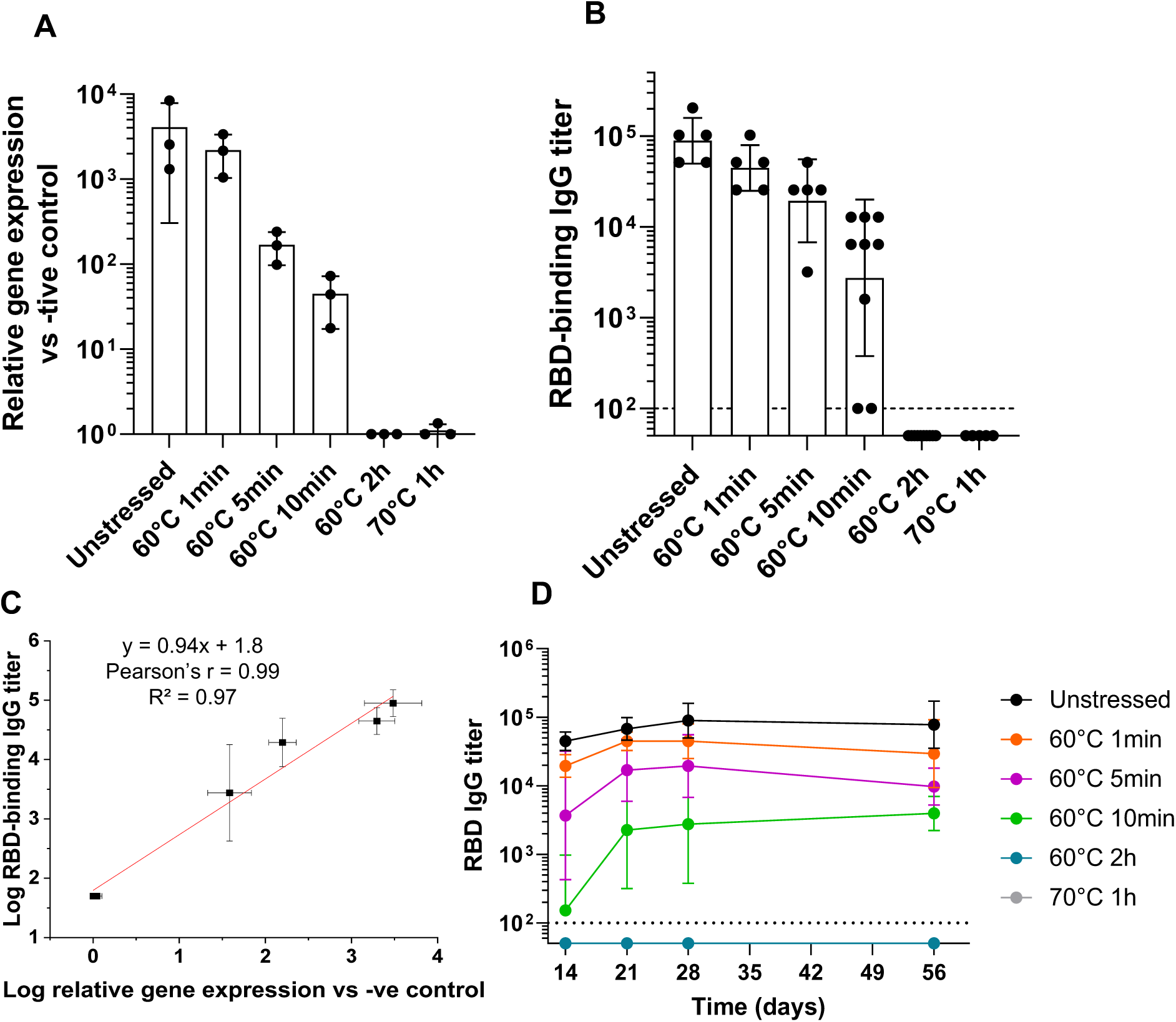
Results of *in vivo* mouse immunogenicity studies for unstressed vs. heat-stressed AC3 samples (A) Relative gene expression levels of heat-stressed AC3 samples (vs unstressed control), that were subsequently dosed in mouse studies, as measured by *in vitro* transduction efficiency. Data are represented as mean ± SD of 3 independent assay replicates. (B) *In vivo* mouse immunogenicity results of the AC3 samples (described in panel A) measured as total antibody titers (vs RBD) in C57BL/6J mice (n≥5) vaccinated with 10E+11 gc of AC3 28 days after vaccination. (C) Correlation of *in vitro* transduction efficiency with *in vivo* potency results shown in panels A and B, respectively. (D) Longitudinal antibody responses in mice (n≥5) treated with 10E+11 gc of AC3 before and after thermal stress. Data are represented as geometric mean ± SD.

## Discussion

### rAAV vectors for gene therapy vs vaccination and associated formulation differences

Recombinant adeno-associated virus (rAAV) vectors are widely studied as gene therapy agents for treating a wide spectrum of human diseases, and there are currently three commercially-available rAAV gene therapy products: Luxturna^®^ (for inherited retinal dystrophy, approved in 2017), Zolgensma^®^ (spinal muscular atrophy, 2019), and Hemgenix^®^ (congenital Factor IX deficiency, 2022). In addition, in Europe, Glybera^®^ indicated for lipoprotein lipase deficiency (now withdrawn by the sponsor due to commercial failure ^51^), Roctavian^®^ and Upstaza^®^ were approved ^52^. There are over 200 ongoing gene therapy clinical trials with rAAV vectors to address unmet medical needs in diverse therapeutic areas such as ophthalmology, neurology, metabolic, hematology and musculoskeletal with over 3000 patients treated over the past 20 years ^53^. From a formulation and delivery perspective, routes of administration include common parenteral sites for systemic delivery (i.e., IV, IM) as well as target-specific locations including the eye, spine, brain, joints, and heart. rAAV vector-based gene therapy products and clinical candidates are commonly stored as frozen liquids formulated in simple PBS-type solutions ^54^, although the recently approved Hemgenix^®^ is stored as a liquid formulation at 2-8°C (the product is formulated in PBS solution with additional stabilizers including sucrose and polysorbate 80) ^55^.

In contrast, despite rAAV viral vectors offering much promise as vaccines, investigations into their utility for this application are more limited. Examples include active immunotherapy strategies (delivery of viral genes to induce protective immune responses) as well as passive immunization approaches (delivery of genes encoding broadly neutralizing antibodies against the virus) in infectious disease targets including HSV, HIV, HPV, influenza and SARS-CoV-2 ^56^. In general, viral vector-based vaccine platforms are a safe, efficacious, low-cost, thermostable approach for genetic vaccination ^57^ as clearly demonstrated with adenovirus vector-based COVID-19 vaccines ^58^. Potential additional advantages for rAAV-based vaccines include its non-pathogenic nature and availability of many different serotypes and capsid variants to develop prime/boost strategies to circumvent anti-vector host antibodies ^59,60^. The rAAV-based COVID vaccine candidate described in this work (AC3) was recently reported to provide long-term immunogenicity and protection in non-human primates ^38,40^. From a vaccine formulation and delivery perspective, rAAV vectors offer the potential for low-cost vaccines including room temperature storage (high inherent capsid thermostability) and non-parenteral administration (oral delivery). Non-parenteral, oral delivery of rAAV vectors not only reduces vaccine administration costs (i.e., eliminating needles, reducing medical waste and less training)^10,23^, but also the potential to enhance global access and improve immune protection (i.e., enhanced mucosal immunity) ^61^. Wildtype AAVs can naturally infect humans via the oral route, are found in human GI secretions and gut tissues ^62^, and are resistant to heat (56 °C for 1 hour) and acidic pH (∼3) exposure ^63,64^, and thus the development of stable, orally-delivered rAAV vector-based vaccines is a realistic formulation goal.

### Physicochemical properties and mechanisms of thermal degradation of rAAV

We employed a wide variety of analytical characterization tools to determine the critical structural attributes (CQAs) of the AC3 rAAV vector. Although recent reviews have highlighted potential key structural properties of rAAV vectors for use as gene therapy candidates^65–67^, no actual data sets with representative rAAV vectors were provided. At the same time, there are numerous recent reports employing individual analytical methods to monitor specific structural attributes of rAAV vectors including chromatography, electrophoresis, light scattering, SV-AUC, TEM, immunoassays and mass spectrometry ^48,68–74^, yet only a few showing a combination of comparative data sets ^72,75,76^. Quality-by-design analysis for rAAV vectors for gene therapy applications have proposed the following CQAs: encapsidated capsids, empty-partially filled capsid ratios, aggregates and deamidated forms ^77,78^. In this work, we report physicochemical characterization and biological potency data with the AC3 vector to determine CQAs from a vaccine development perspective.

Primary structure analysis of AC3 vector showed the expected molecular weight values and capsid composition ratios of VP1, VP2 and VP3 ^70,76^. In addition, post-translational modifications (e.g., truncation, N-terminal acetylation) in AC3 were overall similar to reports of product-related species with other rAAV vectors ^79,80^. Importantly, relatively low levels of deamidation were observed at N57 (∼25%) and N510 (∼5%), which are highly conserved across AAV serotypes and Asn deamidation at these sites can impact transduction efficiency and T cell response, respectively ^81,82^. Since conventional trypsin- or chymotrypsin-based proteolytic methods can induce substantial artificial deamidation in rAAVs during sample handling ^72,83^, we utilized the speed and efficiency of a recently commercialized kit (S-TrapTM) to rapidly generate LCMS-compatible tryptic or chymotryptic peptides of the VP proteins in AC3 to limit assay-related deamidation ^84^.

Particle size analyses of the AC3 vector demonstrated a relatively homogenous viral capsid population (∼30 nm diameter), in agreement with rAAV size range reported other studies ^85,86^, with no notable aggregate formation. The percent full-capsids (∼36%) in the AC3 preparation was determined by TEM and SV-AUC, however, TEM analysis cannot currently distinguish empty vs. partially-filled capsids ^87^. SV-AUC is thus the current “gold standard” to evaluate the relative proportions of full, partially-filled and empty capsids as well as aggregates ^64,88,89^, yet this is a highly specialized technique, so efforts to identify alternative, more easily implemented analytical approaches (e.g., chromatographic, electrophoretic and mass spectrometry) are ongoing ^76^. Such compositional analyses of rAAV preparations are performed to monitor lot-to-lot consistency, yet reports of their effects on gene transduction or *in vivo* performance are limited ^64,90^. From a rAAV vaccine perspective, a possible correlation between a higher full/empty capsid ratio with heightened immune responses was reported ^91^.

Thermal degradation studies with rAAV vectors have primarily focused on batch-to-batch consistency evaluations ^48,92^ or rAAV serotype identity testing ^49^ by utilizing extrinsic fluorescence (e.g., differential scanning fluorimetry, DSF). In this work, we employed DSF along with light scattering, intrinsic fluorescence spectroscopy and SV-AUC analyses to elucidate the AC3 vector’s molecular mechanisms of thermal degradation. Three structural transitions were observed for AC3 vector during thermal stress: (1) a “small” region where capsid proteins undergo subtle structural transitions between 35-55°C, (2) a “medium” region between 55-65°C where the amount of uncoated genome increases steadily without capsid disassembly, and finally (3) a “major” region between 64-70°C where capsid disassembly and genome release both rapidly occur. Our results can be evaluated in the context of previously proposed rAAV degradation mechanisms including ssDNA genome translocation (or ejection) from intact capsids, and/or rAAV capsid disassembly due to protein denaturation. ^46,93^ For example, SV-AUC analysis of thermally stressed AC3 samples showed a decrease in full (and partially-filled) capsids with a concomitant increase in both empty capsids and small molecular weight impurities. Such results are consistent a capsid disassembly-based degradation mechanism at higher temperatures (“major transition”), since a proportional conversion of full to empty capsids was not observed^47^. At the same time, results from *in vitro* gene transduction assays of thermal stressed AC3 samples (see next section) highlight a link to “minor” and “medium” structural transitions. Interestingly, recent reports examining rAAV degradation during freeze-thaw (F-T) stress ^46,47,92^ have observed genome leakage in the absence of major structural alterations to the capsid, suggesting rAAV freeze-thaw degradation mechanism follows a genome translocation-based model. Interestingly, excipients can either enhance rAAV stability against F-T stress (e.g., surfactants and cryoprotectants) or cause further instability (e.g., due to excipient crystallization during sample thawing)^46^. Such excipient evaluations with AC3 during thermal stress will be of interest for future work to identify optimized heat-stable formulations. In addition, for oral delivery, stabilizers to prevent vector degradation *in vivo* (by gastric acid and digestive enzymes) will need to be identified as future work, as has been reported with orally administered live rotavirus vaccines ^50^.

### Development of cell-based in vitro gene expression assay (RT-qPCR) and correlations with in vivo immunogenicity

To dose rAAV vectors for gene therapy applications, genome copy number (e.g., total and/or within infectious capsids) are routinely determined ^94,95^ typically by employing PCR-based methods ^96^. Such approaches, however, are not indicative of the number of fully active viral particles that dictate gene expression levels or related biological efficacy readouts (e.g., gene augmentation or gene silencing). To this end, rAAV potency assays are typically required including *in vitro* cell-based assays or *in vivo* animal models. For example, Clarner et al developed a one-step RT-ddPCR method to quantify the potency of different rAAV vectors (for gene augmentation or repression applications) by determining target mRNA levels in HeLa cells (*in vitro*) and correlating results to expression levels in non-human primate (*in vivo*) samples ^97^. Cell-based potency assays are preferred for facilitating process and formulation development work, since animal models are expensive, low-throughput and time-consuming. Nonetheless, Gruntman et al reported measuring transgene expression levels in mice to determine rAAV vector stability under conditions encountered in gene therapy clinical trials ^98^.

For dosing of vaccine candidates, either mass (e.g., protein concentration assays for recombinant proteins and non-replicating viral vectors) or titers (e.g., viral infectivity assays for live viruses) are typically measured. At the same time, the ability to generate an immune response *in vivo* is the key readout for developing an informative potency assay including either animal models or surrogate *in vitro* cell-based assays ^20,99^. Similar to the animal models described above for gene-therapy, *in vivo* immunogenicity assays are also time-consuming and expensive, and their replacement with *in vitro* based readouts of immunogenicity is also a high priority as part of the 3Rs principles of replacement, reduction, and refinement for animal use in research ^100^. In the case of genetic vaccines (e.g., viral vectors or mRNA-LNPs), measuring gene expression levels in a cell-based assay is a reasonable approach if correlations with immune responses *in vivo* can be established.

In this work, we developed an *in vitro* cell-based gene expression assay for the AC3 vector and then correlated the results with *in vivo* immunogenicity levels in mice. This gene transduction efficiency method detects mature AC3 mRNA using a specific TaqMan™ probe in a two-step RT-qPCR assay using Huh7 cells. We utilized a recently published improved method by Sanmiguel et al to accurately quantify the rAAV genomes with or without DNase-I treatment (37°C for 1 h) ^101^. We established linear range, selectivity and stability-indication, the latter by using thermally stressed AC3 samples, and a strong correlation (Pearson’s r -0.93 and -0.94) between gene transduction efficiency and *in vivo* mouse immunogenicity results were observed.

For the animal model potency assay, rAAV vector AC3 can induce a durable humoral and T-cell responses with a single dose in mice, potentially requiring none or fewer boosters compared to other vaccine platforms ^38^. In this work, high levels of measured RBD IgG titers, that reached maximum levels at day 28 with no decline up to the last studied data point (day 56), were observed. Recent studies have demonstrated a single dose of AC3 to be immunogenic in mouse and non-human primates and the peak neutralizing antibody titers remained unchanged up to the length of the 20 months in NHP which was complemented by cellular immunity ^38,40^. Although more extensive studies in animals and humans will be required to determine safety, efficacy, and durability of these immune responses, these results suggest that total antibody titers in mice, instead of neutralizing antibody titers, may be sufficient for use as an analytical potency assay, especially in the context of establishing a surrogate *in vitro* cell-based potency assay.

### Correlations of physically stressed AC3 samples with the cell-based in vitro transduction assay and in vivo mouse immunogenicity studies

We demonstrated a high degree of concordance between relative gene expression levels of AC3 vector samples (unstressed and stressed) as measured by *in vitro* RT-qPCR based method and total antibody titers in an *in vivo* mouse immunogenicity assay. Although *in vivo* immunogenicity assays commonly display a linear range with varying antigen doses, it is not necessarily the case that stressed antigen samples demonstrate such behavior. For example, many traditional vaccine antigen platforms (e.g., inactivated viruses and recombinant proteins) can undergo significant loss in their antigenicity values as measured by *in vitro* potency assays, but *in vivo* assays are more forgiving and a relatively small loss in immunogenicity is observed. Recent examples of this phenomenon include stressed samples of inactivated polio vaccine ^102^ and a SARS-CoV-2 RBD vaccine candidate ^103^. In this work, however, AC3 samples with increasing exposure to thermal stress correlated with reduced gene expression (*in vitro* cell-based assay) and total antibody (mouse immunogenicity assay) levels. Presumably, this proportional effect with stressed rAAV samples is due to differences in the vaccine modalities since genetic vaccination requires transgene expression to produce the antigen vs. traditional vaccines where administering stressed antigen may not completely abrogate all of the immunogenic epitopes across the dose range in an animal model.

The same thermally stressed AC3 samples that showed correlations between the *in vitro* and *in vivo* potency assays were also examined by physicochemical analyses. Interestingly, a strong correlation in the readout of some of these methods was observed, but not for others. For example, a direct correlation was observed with genome copy levels (after DNase treatment) for stressed samples showing encapsidated gene levels (not total gene copies) are an important stability-indicating parameter for AC3. In addition, inverse correlations were observed with gene expression levels and increasing levels of SYBR gold fluorescence signal (i.e., loss of encapsidated genomes) and DLS signal (aggregate formation). No such correlations were observed with major structural alterations of the capsid (i.e., thermal melting temperatures or subvisible particle formation) indicating the loss of potency had occurred at temperatures below where such dramatic alterations to capsid structure had occurred. We also examined the *in vitro* gene expression levels of AC3 samples stressed during thermal ramp studies using intrinsic and extrinsic fluorescence spectroscopy. No loss in relative gene expression was observed during the thermal ramp up to 50°C, with a notable loss observed at 60°C, and complete losses observed by 70°C. Similar trends were observed for encapsidated genome copies as measured by the genome copy assay.

These biophysical data sets demonstrate that temperature exposure below 50°C can cause subtle alterations in the AC3 capsid structure which do not affect potency. When exposed to temperatures in the range of 55-65°C, AC3 capsid shows some structural alterations and leakage of DNA from capsid which correlate with partial loss of *in vitro* potency. Finally, at higher temperatures of 70°C, *in vitro* potency is lost when the capsid extensively degrades. In summary, these stability results with the AC3 vaccine vector indicate gene expression levels in cells, immunogenicity levels in mice, and the number of encapsidated genomes are linked in a stressed AC3 sample.

### Conclusions and future challenges to formulate rAAV vectors as vaccine candidates

In this work, we employed an extensive analytical toolbox to characterize a second generation, rAAV vector-based COVID-19 vaccine candidate (AC3). The AC3 vector was analyzed in terms of key structural attributes including primary structure/PTMs of the VP proteins, viral capsid assembly (e.g., size, ratio of empty to partially-filled to full capsids, aggregates), gene copies (both total and encaspidated), gene expression in a cell-based RT-qPCR assay and *in vivo* immunogenicity in mice. Moreover, we report the inter-relationships between the biological activity and physicochemical properties and stability of AC3 vector by evaluating different stressed samples in terms capsid degradation profiles, gene transduction efficiency and *in vivo* performance.

By identifying stability-indicating methods and elucidating the mechanisms of AC3 vector degradation, these results establish critical quality attribute (CQAs) of rAAV for use as a viral vector vaccine, and thus support future Chemistry, Manufacturing and Control (CMC) activities. For example, during process scale-up, analytical comparability studies will be required to demonstrate similarity between rAAV vectors produced from new and old processes ^104^. By employing the physical stability-indicating assays identified in this work (e.g., changes in encapsidated genomes) as well as *in vitro* potency assays (e.g., gene transduction efficiency assay in cell-culture), the screening of stabilizing excipients can be performed to identify rAAV-based vaccine formulations. For example, orally administered, liquid formulations of rAAV vectors that are refrigerator (or ambient temperature) stable have the potential to lower costs, improve accessibility and increase global vaccine coverage especially in LMICs ^105,106^.

## Limitations of the study and future directions

Additional studies are required to assess the general applicability of these findings with other rAAV-based vaccine candidates, especially in terms of the inter-relationships between physiochemical properties and biological potency, especially *in vivo* immunogenicity. Evaluation of how different capsid species observed to be present in purified, unstressed AC3 samples (i.e., partially-filled and deamidated capsids) may affect *in vitro* transduction efficiency and *in vivo* immunogenicity is suggested. Finally, possible interference of excipients required to stabilize rAAV vectors with the physicochemical and cell-based analytical tools employed in this study will need to be assessed.

## Materials and Methods

### Materials

The primers and probes for *in vitro* potency assay and genome copy assay were received from Massachusetts Eye and Ear Institute. TaqMan™ Universal PCR Master Mix, no AmpErase™ UNG, high-capacity cDNA reverse transcriptase kit, and PCR buffer were purchased form Applied Biosystems. DNase-I was procured from Sigma-Aldrich. All reagents and chemicals used were of analytical grade and purchased from Sigma-Aldrich.

Purified AC3 vector stock was produced by Novartis Gene Therapies as previously described (PMID: 34428428). AC3 stock solution was obtained at 8.83E+12 gc/mL in buffer (20 mM Tris buffer containing 1 mM MgCl2, 200 mM NaCl, 0.005% Pluronic F68, pH 8.1) and was stored at -80°C before use. The viral stock solution was diluted 10-fold corresponding to 8.83E+11 gc/mL in buffer (20 mM HEPES buffer containing 1 mM MgCl_2_, 200 mM NaCl, 0.005% Pluronic F68, pH 7.0), unless otherwise specified in the text. The diluted AC3 samples, unstressed or after thermal stress were examined using various analytical techniques as described below.

### Methods

#### SDS-PAGE

SDS-PAGE was performed as adapted from Agrawal *et al*, 2020 ^107^. AC3 stock diluted to 9.81E+10 gc/mL was loaded into the SDS-PAGE sample mixture. 5 uL of Novex Sharp Pre-Stained Protein Standard molecular weight ladder (Invitrogen) were loaded alongside the samples. Samples were separated at 120V for 10 min, followed by 150V for 50 minutes. Protein bands were visualized with a Pierce Silver Stain kit (Thermo-Fisher) and quantified using AlphaView gel imaging software.

#### CE-SDS

CE-SDS was performed following an adapted procedure from Zhang & Meagher ^70^. For reducing conditions, AC3 was treated with 150 mg/mL SDS and 10% v/v β-Me, and incubated for 5 min at 95°C. For non-reducing conditions, AC3 was treated with 150 mg/mL SDS and 60 mM iodoacetamide, and incubated for 30 min at 60°C. The reduced or non-reduced AC3 samples were then buffer-exchanged into 0.5 mg/mL SDS with or without 5% v/v β-Me, respectively. Finally, a 10 kDa MW internal standard (SDS-MW kit, AB SciEx LLC) was added. The AC3 samples were electrokinetically injected (-5kV for 60 sec) into a bare fused capillary (50 µm I.D., 40 cm effective length; Agilent Technologies) using a 7100 CE system (Agilent Technologies). AC3 capsid proteins (viral proteins VP1-3, see **Figure 1**) separation was achieved using a -28 kV over 45 min and were detected using a wavelength of 214 nm. AC3 electropherograms were integrated using MassHunter Qualitative Analysis v10.0 software (Agilent Technologies).

#### Intact Mass Analysis

Prior to intact mass analysis, AC3 was treated with 10% acetic acid and incubated for 15 min at RT ^79^. The sample was centrifuged for 5 min at 17,000 x g and the top 90% of sample was transferred to an UHPLC vial for analysis. Approximately 3 µg of AC3 was injected into 1290 Infinity II UHPLC system (Agilent Technologies) containing a Premier CSH-C18 column (Waters Corporation). The column and autosampler compartment temperatures were 80 and 10°C, respectively. The LC mobile phases consisted of water with 0.1% formic acid (mobile phase A) and acetonitrile with 0.1% formic acid (mobile phase B). VP (see **Figure 1**) protein separation was achieved using a 20 min 20-40% mobile phase B gradient, and VP elution was monitored using a wavelength of 214 nm. VP protein masses were then measured using an in-line 6545XT QTOF system (Agilent Technologies) using the following parameters: 4000V VCap, 180V fragmentor, 290°C gas temperature, 600-2200 m/z scan range per sec, and masses were corrected using a 922.0098 m/z reference mass. Mass spectra were deconvoluted using MassHunter Bioconfirm v10.0 software (Agilent Technologies).

#### LC-MS peptide mapping

Prior to LCMS analysis, VP1-3 peptides were generated using a commercial kit and associated protocol (S-Trap^TM^ micro kit, Protifi LLC). Briefly, approximately 5 µg of AC3 was reduced with TCEP, heat denatured (15 min at 55°C), and then alkylated with methyl methanethiolsulfonate (MMTS). VP1-3 proteins were then bound to the S-Trap column and digested with trypsin (Promega Corporation) for 2 hrs at 47°C. VP peptides were then analyzed using the same LCMS system as described for Intact Mass Analysis. VP peptides were injected onto an Advanced Peptide column (2.1 x 150 mm, 2.7 µm, Agilent Technologies), and separated using a LC gradient of 2-45% B (A: 0.1% formic acid in water, B: 0.1% formic acid in acetonitrile) over 74 min at a flow rate of 0.3 mL/min.

The electrospray ionization parameters consisted of: 325°C gas temperature, 4000V Vcap, and 100V fragmentor. Mass spectra were collected from 250-1700 m/z at 1 spectra/sec. The threshold for MS/MS analysis was 40,000 counts and the two most abundant ions were selected for CID fragmentation per cycle. Mass spectra were processed using MassHunter Bioconfirm v10.0 software (Agilent Technologies), with the following variable modifications included: Cys alkylation, Met oxidation, Asn deamidation, N-terminal acetylation, and Ser/Thr/Tyr phosphorylation.

#### DLS

Twenty-five µL of AC3 was loaded at 8.83E+11 gc/mL in each well of a 384 well plate, centrifuged briefly, and DLS was performed using a DynaPro Plate Reader III (Wyatt, Technology, CA). Five separate acquisitions were taken for 5 s each and averaged, and this procedure was repeated in triplicate. Data were fit to lognormal (i.e., cumulant) and mean squared displacement (i.e., multimodal) distributions and plotted as a function of intensity. Data were corrected for solvent viscosity and all measurements were recorded using a thermal ramp from 20-80°C in 1°C increments.

#### TEM

Five µL of AC3 was adsorbed to a carbon coated copper grid for 2 minutes (Ted Pella), remaining sample wicked out, and negatively stained with 2% (w/v) Uranyl acetate for 3 minutes. The dye was removed by using an adsorbent paper without a wash step and air dried before acquiring images in 200kV acceleration voltage. Particle size of AC3 was calculated using Image J software ^108^ and percentage full capsids were calculated by manual counting of 250 particles as described in detail in the results sections.

#### SV-AUC

SV-AUC was performed following the method of Burnham et al 2015 ^88^. Briefly, the AC3 samples were diluted to a final OD260 (0.2), which corresponds to ∼ 2E+12 gc/mL in 20mM Tris, 1mM MgCl_2_, 200mM NaCl, 0.005% PF-68, pH 8.1 and 0.4 mL sample was loaded into Beckman charcoal-epon two sector cell with 12 mm centerpieces and either sapphire or quartz windows. Sedimentation velocity experiments were performed using an Optima analytical ultracentrifuge equipped with a scanning ultraviolet-visible optical system (Beckman Coulter, Indianapolis, IN.) All experiments were performed at 20°C after at least 1 h of equilibration after the rotor reached 20°C. A rotor speed of 20,000 rpm, UV detection at 260 nm, and a scan frequency of 60 s were used for a total of 150 scans. The data were analyzed using Sedfit (Peter Schuck, NIH) fitting the data to a continuous c(s) distribution from 0 to 200 svedbergs. A resolution of 300 points per distribution and a confidence level of 0.95 were applied to all fits. Baseline, radial independent noise, and time independent noise were fit, while the meniscus and bottom positions were set manually. The c(s) distributions were imported into Origin 2018 (OriginLab, Northampton, MA) for peak integrations and graph generation.

#### IF and SLS

AC3 was diluted to 50 µg/mL in 20mM Tris, 1mM MgCl_2_, 200mM NaCl, 0.005% PF-68, pH 8 in 0.2 x 1 cm quartz cuvettes in 250 µL final volume. Fluorescence was measured using a QM-40 Spectrofluorometer (Horiba) equipped with a 4-position cell holder, a peltier temperature control device, a 75W Xe arc lamp, and a Hamamatsu R1527 photomultiplier tube. Fluorescence emission spectra were recorded from 10-87.5°C in 2.5°C increments, 1°C/min ramp rate, and 2 min hold time at each temperature. Three samples and one buffer blank were measured using an excitation wavelength of 295 nm (>95% for Trp emission) for each cuvette set up. Emission spectra were collected from 310-390 nm with a step size of 1 nm and an integration time of 1 s. The excitation and emission slits were both set at 4 nm. The buffer emission was subtracted from the sample emission and the MSM peak fitting algorithm was executed in Origin 2018. Light scattering at the excitation wavelength (295 nm) was recorded using a second photomultiplier 180°C from the fluorescence detector and was plotted as a function of increasing temperature.

#### DSF

The fluorescent dyes (SYPRO Orange or SYBR Gold) were individually diluted to a final concentration of 50X in dPBS. In each PCR well, 45 µL of diluted AC3 (8.83E+11 gc/mL) or buffer and 5 µL of diluted florescent dye was added and mixed well by pipetting up and down a few times. Fluorescence emission spectra were recorded from 25-92°C in 1°C increments, and 2 min hold time at each temperature using Stratagene DSF Instrument (Agilent, Model 401513). Fluorescence data was collected using three samples and three buffer blanks using SYPRO (610 nm) and FAM (516 nm) filters for SYPRO Orange and SYBR Gold dyes, respectively. The buffer emission was subtracted from the sample emission and the results were plotted as (1) fluorescent intensity vs temperature, and (2) as fluorescent intensity of unstressed sample at 25°C.

#### Thermal stability studies

For accelerated stability studies, 500 µL of diluted AC3 samples in 1.5 mL polypropylene tubes were subjected to incubation at 60°C (from 1 min up to 2 hrs.) or 70°C (1 hr.) in a block heater and compared to unstressed control (4°C) samples. For mechanistic studies, AC3 samples were loaded into plate readers/cuvettes and subjected to thermal ramps as described earlier for individual analytical techniques (Intrinsic fluorescence (IF), DSF using SYPRO orange and SYBR Gold dyes, DLS and SLS). Buffer alone samples were also run and subtracted from AC3 data sets from each instrument. Additionally, AC3 was also stressed using the DSF SYBR Gold method in absence of a dye and effect of thermal ramp treatment was studied using genome titration and transduction efficiency assays as described below.

#### Genome titration assay

Genome titration assay of AC3 was performed using a two-step method as described by Sanmiguel *et al*, 2019 ^101^. Briefly, AC3 was first treated with/without DNase-I (37°C 1h) followed by 10^5^ folds serial dilution. This diluted AC3 was then subjected to droplet digital Polymerase Chain Reaction (ddPCR) assay using TaqMan™ universal master mix, primers (forward: GTGCAGCCAACCGAG, reverse: ACACCTCGCCAAATGG), probe (6FAM-TCTATCGTGCGCTTTC-MGBNFQ) under cycling thermal conditions (step 1: 95°C for 10 min; step 2: 40 cycles of 95°C, 15 s and 60°C, 1 min, data collection at 60°C) using the Bio-Rad’s QX200 Droplet ddPCR System system. Final titers (gc/mL) were calculated using the following equation:

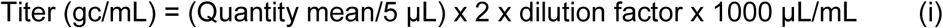

#### Transduction efficiency assay

Transduction efficiency (also referred as *in vitro* infectivity or *in vitro* potency assay) of AC3 was performed using a two-step real-time quantitative Polymerase Chain Reaction (RT-qPCR) assay (workflow presented in **Supplemental Figure S1**). 80-90% confluent Huh7 cells monolayers in 96-well plates were sequentially infected with required dilutions of helper virus (wildtype human adenovirus 5, wtAdeno5, used at multiplicity of infection (MOI) of 20) and AC3 test articles (diluted to a 2E+8 gc/well, resulting in an estimated MOI of 1E+4) and negative control (wt Adeno 5 only control, no AC3) followed by 72 h incubation at 37°C, 5% CO_2_. Total RNA was extracted by adding chilled buffer (10mM Tris-HCl pH 7.5, 150mM NaCl, 0.1% Igepal CA-630), and incubating at -80°C for 1h followed by supernatant transfer to PCR 96-well plate and kept chilled or at -80°C until used. The reverse-transcription (RT) step was performed using MultiScribe^TM^ Reverse Transcriptase and random primers under thermal cycling conditions (step 1: 25°C for 10 min, Step 2: 37°C for 120 min, step 3: 85°C for 5 min, step 4: 4°C for ∞) adapted from the manufacturer ^109^, using QuantStudio^TM^ 7 Flex Real-Time PCR System (Applied Biosystems, USA). The quantitative Polymerase Chain Reaction (qPCR) step was performed using TaqMan™ universal master mix, primers (Forward: CAGATCCTGCAGAAGTTGG, reverse: AGCAGCACACAGCAG), probe (6FAM-TGGGCAGGTGTCCAG-MGBNFQ) under cycling thermal conditions (step 1: 95°C for 10 min; step 2: 40 cycles of 95°C, 15 s and 60°C, 1 min, data collection at 60°C) using the QuantStudio^TM^ 7 system. Quantification of gene expression was reported relative to negative control calculated using the following equations, where Ct denotes threshold cycle.

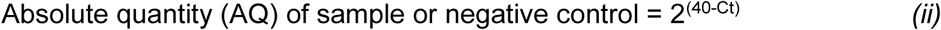

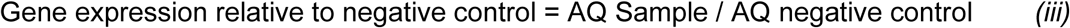

### In vivo mouse immunogenicity assay

Mouse studies were performed in compliance with the Schepens Eye Research Institute IACUC. The C57BL/6 female mice were purchased from Jackson Laboratories. Animals were housed in standard BSL1 facilities, with 12-hour light cycles and free access to regular chow diet and water. 6–9-week-old mice were intramuscularly (right gastrocnemius muscle) treated with 10^11^ gc/mouse of unstressed or stressed AC3 vaccine. Serum samples were obtained by submandibular bleeds for humoral immune response analyses. RBD binding antibodies titers were determined using a published protocol ^38^. Briefly, ELISA plates coated with 1 mg/ml SARS-CoV-2 RBD were blocked with blocker casein in PBS followed by addition of serially diluted serum samples and incubated for 1h at RT. The plate were then loaded with secondary rabbit anti-mouse IgG for 1h at RT followed by color development using TMB and read at 450 nm.

## Data availability statement

The dataset presented in the current study are available in the KU ScholarWorks repository (to be updated). The data is also available with the corresponding authors. Supplementary materials associated with this article can be found in the online version at doi (to be updated).

## Acknowledgements

The authors would like to thank Saleh-Birdjandi Soraia from Vaccine Analytics and Formulation Center at University of Kansas for technical assistance.

## Author contributions

Planning, conducting experiments and manuscript preparation: PK, MW, OSK, JH, NZ, JS. Supervision, scientific guidance, planning experiments and reviewing and editing manuscript: DBV, SBJ, LHV. All authors read and approved final version of the manuscript.

