## Supplemental Information for "Correlating physicochemical and biological properties to define critical quality attributes of a recombinant AAV vaccine candidate"

**Supplemental Methods and Figures**

*Supplemental Methods*

**Subvisible particles:** Analysis of subvisible particles was performed using the Horizon Sub-visible Particle Analysis Instrument (Halo Labs, CA). Unstressed and thermally stressed AAV AC3 samples were run at a 10-fold dilution. A volume of 25 µL of each sample was transferred to a special Halo Labs 96-well plate and analyzed in triplicate on the Horizon Sub-Visible Particle Analysis Instrument. Sample measurements were buffer-subtracted and the system was programmed to detect subvisible particles in the size range between 2-100 um in diameter.

**Supplemental Figure S1**: Workflow and validation of *in vitro* transduction efficiency assay.

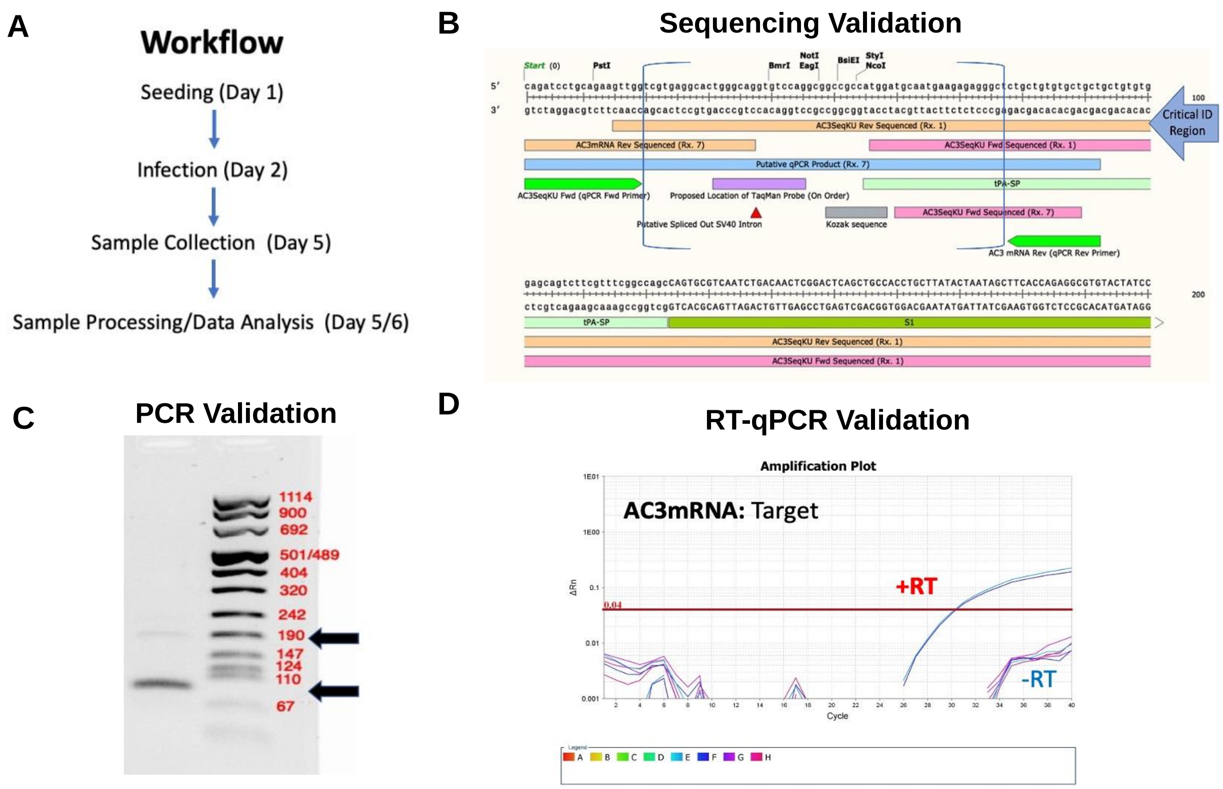

**Supplemental Figure S2**: Primary sequence coverage of AAV AC3 using LCMS peptide mapping. Sequence coverage from chymotrypsin- (red bars) or trypsin-digestion (black bars). The first residue of VP1, VP2, and VP3 proteins, and the location of the VP3 truncations (denoted with an asterisk) are indicated. The combined trypsin and chymotrypsin peptide maps covered ~98% of the VP1 primary sequence.

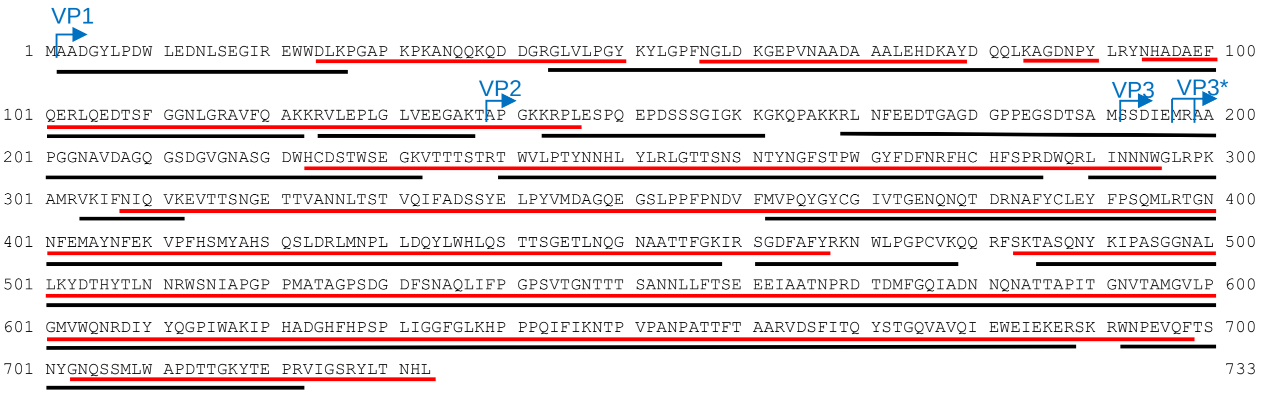

**Supplemental Figure S3:** Additional biophysical method measurements of unstressed vs heat-stressed AC3 samples that did not correlate well with relative gene expression levels obtained from *in vitro* transduction efficiency assay. Correlation plots between AAV AC3 relative gene expression levels versus (A) sub-visible particles obtained from HALO analysis, (B) Tm values from DSF SYBR Gold analysis vs temperature, (C) fluorescence intensity at 25°C from SYPRO orange fluorescence, and (D) Tm values from SYPRO Orange fluorescence vs temperature. Mid-point transition temperature (Tm) was calculated using 1st derivative function in Origin Lab Software (Origin 2018).

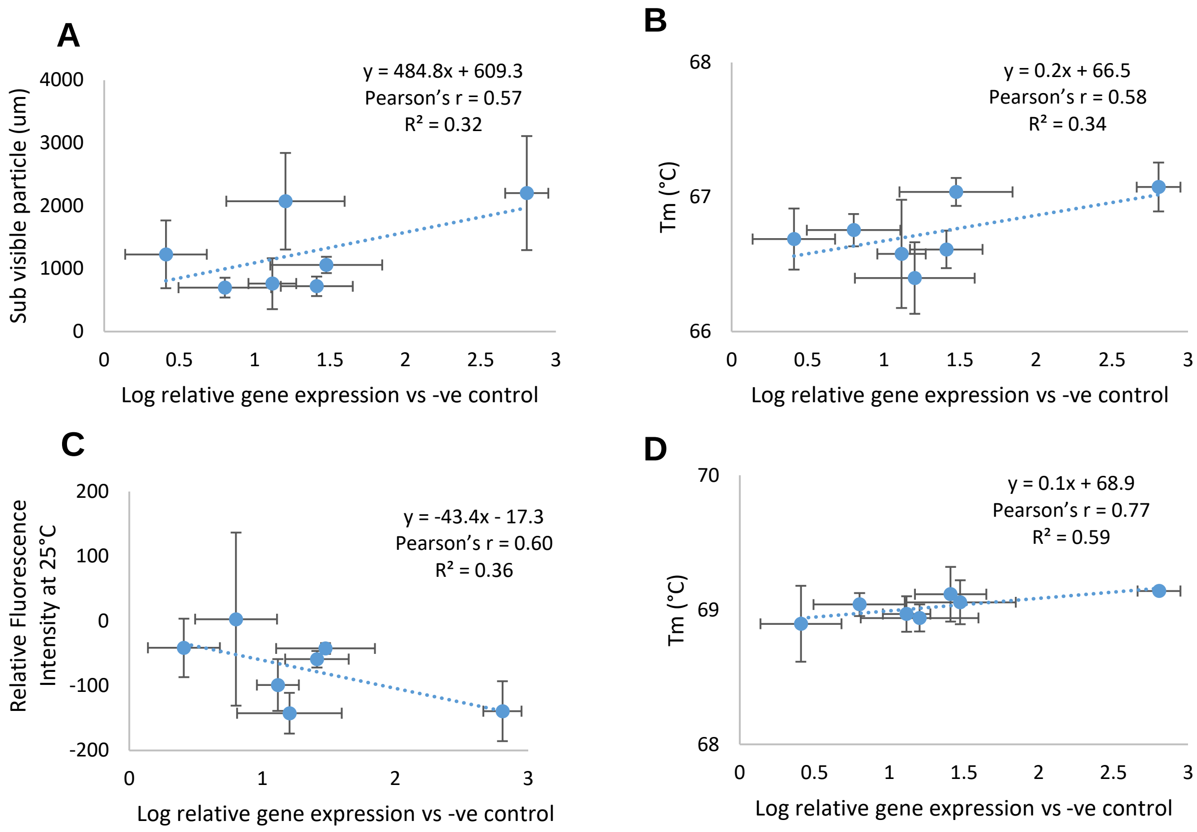

**Supplemental Figure S4:** Sub-visible particle analysis of (A) unstressed and (B) thermal-stressed (10 min at 60°C) AC3 vector as measured by Horizon Sub-visible Particle Analysis Instrument.

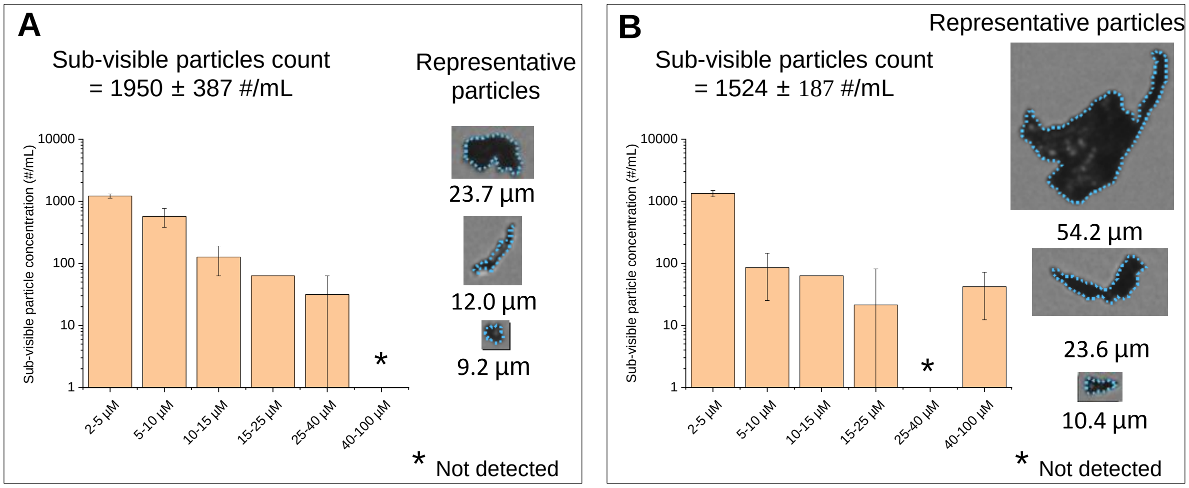
